# Hierarchy in the mouse frontal cortex in mnemonic olfactory decision-making

**DOI:** 10.64898/2026.05.20.726442

**Authors:** Josefine Reuschenbach, Maria Inês Ribeiro, Xiaochen Fu, Izumi Fukunaga

## Abstract

The prefrontal cortex plays a critical role in integrating the memory of a recent experience to guide context-dependent decisions, yet how the finer, sub-steps of decision formation are physiologically implemented remains poorly understood. Using an olfactory delayed non-match-to-sample task with graded stimulus similarity in head-fixed mice, we examined how decisions emerge when a current sensory event and a memory of a recent event must be compared. With high-density extracellular recordings across the frontal cortex, we characterized odor-specific delay activity and decision-related signals. Interestingly, the secondary motor cortex showed minimal sensory coding, suggesting that areas engaged in mnemonic decision-making differ from those involved in simpler, stimulus-response decision-making in the rodent brain. Rather, there was a gradual emergence from sensory representations to choice-related activity, with intermediate regions showing choice modulations that retain stimulus sensitivity that arises with early timing, as well as balanced match vs. non-match selectivity. These results suggest that mnemonic decision-making is supported by a distributed frontal network in which sensory and choice signals are gradually integrated, rather than localized to a single comparator region.

## Introduction

Animals in their natural environment need to rely on the memories of recent events to adjust their behavior. This is evident in food seeking behavior, where animals must avoid repeatedly foraging in recently depleted patches (Applegate & Aronov, 2022; Owen-Smith *et al*, 2010; Rudebeck & Izquierdo, 2022). The prefrontal cortex (PFC) is considered essential for such mnemonic behavior (Fuster, 1988; Goldman-Rakic, 1995; Hanks *et al*, 2015; Kim & Shadlen, 1999; Le Merre *et al*, 2021; Mante *et al*, 2013; Miller & Cohen, 2001; Passingham, 2021; Romo *et al*, 1999; Siegel *et al*, 2015), but if and how the individual steps involved in such decision-making are physiologically implemented, and where within this interconnected brain region these operations are positioned remain elusive. This gap in the knowledge is especially acute for chemical senses, despite their evolutionary importance and dominance in rodent perception.

Decision-making involving working memory behavior has been modelled experimentally with sensory sequence tasks, including the delayed match to sample task and the delayed non-match-to-sample (DNMS) task. The cognitive elements involved in such tasks include the recognition of specific sensory stimuli and its short-term retention during a delay period, and comparison of past vs. current events (match/non-match computation), culminating in a choice and motor plan (Brody *et al*, 2002; Engel & Wang, 2011; Wu *et al*, 2020). Whereas the initial comparison of the events needs to occur in a stimulus-specific circuit manner, animals easily generalize the task rule over many sensory stimuli (Miller & Cohen, 2001; Nakayama *et al*, 2022; Reuschenbach *et al*, 2023; Wang *et al*, 2026). This suggests that the process involves several steps of operations, involving a stimulus-specific comparison of past and current sensory stimuli, and then an integration of these signals to generate a stimulus-invariant decision variable (Engel & Wang, 2011; Moody *et al*, 1998). It is unclear where exactly these operations occur, and if they correspond to known anatomical divisions in the rodent PFC.

In the context of the olfactory DNMS task in the mouse, the anterolateral motor cortex (ALM) of the frontal lobe, a sub-region within the secondary motor cortex (MOs), has been proposed to compute whether past and recent sensory events match (Wu *et al*., 2020). This is because the neuronal activity in this region reflected all cognitive elements listed above and is sensitive to optogenetic manipulations. However, abundant sensory-driven delay activity is present in the medial PFC and orbitofrontal cortex, at least during the task acquisition phase (Liu *et al*, 2014; Ramus & Eichenbaum, 2000; Wu *et al*., 2020). In addition, there exist ample reciprocal connections between MOs and the medial and ventral prefrontal regions (Harris *et al*, 2019; Heidbreder & Groenewegen, 2003; Hoover & Vertes, 2007; Zingg *et al*, 2014), suggesting that the result of network interactions within PFC may culminate in the output activity of MOs neurons. Thus, while the secondary motor cortex is currently considered the most likely locus of match vs. non-match computation in the rodent brain, it is unclear if this region alone contains most of the steps of match vs. non-match computations, or where, along a potential hierarchy of context-dependent decision making, this region is positioned. Answering this question requires direct comparisons of the frontal cortex subregions in a task that dissociates the sensory elements from decision components.

Here, we combined an olfactory DNMS task with graded odor similarities and dense electrophysiological sampling across the mouse frontal cortex to map where the sensory memory, mnemonic comparison, and abstract match vs. non-match signals emerge. We found that the secondary motor area carries only weak sensory information and primarily value-related, decision variable. In contrast, anatomically intermediate areas, including the prelimbic cortex, exhibit strong sensory coding during delay and early, stimulus-specific match vs. non-match signals. These distinctions may correspond to a progressive generation of mnemonic decision-making along a ventromedial-to-dorsolateral gradient.

## Results

### Mice generalize a DNMS rule across odor pairs with graded similarity

We first established a behavioral paradigm for match vs. non-match decision making that can also resolve fine stimulus-specific information. We used an olfactory DNMS task (Fig. 1) where the mice judged whether a brief sample odor was followed, after a short delay, by the same or a different test odor (match or non-match, respectively). The task was implemented as a Go/No-Go paradigm, where only non-match trials were rewarded. The odor sets comprised four non-matching and four matching odor pairs ranging in perceptual similarity (Reuschenbach *et al*., 2023), including structurally unrelated monomolecular odorants, closely related esters, and pairs differing only in concentration (Fig. 1A,B). We term this paradigm the multiple-odor DNMS task. This design allowed us to vary discrimination difficulty in order to distinguish between odor-specific vs. odor-invariant signals.

Head-fixed mice were first trained on a two-odor version of the DNMS task to acquire the task rule (Methods), and once proficient, they transitioned to the multiple-odor DNMS task (Supplementary Fig. 1). The mice showed high accuracy readily after this transition, generating anticipatory licks across non-matching pairs while withholding anticipatory licking on matching trials (Fig. 1C,D, Supplementary Fig. 1; overall accuracy (auROC)=0.869 +/-0.065, p = 4.01e^-5^ Wilcoxon sign rank test for median = 0.5, n = 22 mice). This indicated that animals could flexibly apply the learned task rule to diverse odor combinations. Thus, this multiple-odor DNMS paradigm provides a robust behavioral assay to interrogate match vs. non-match recognition for odors that vary systematically in similarity.

**Figure 1:**
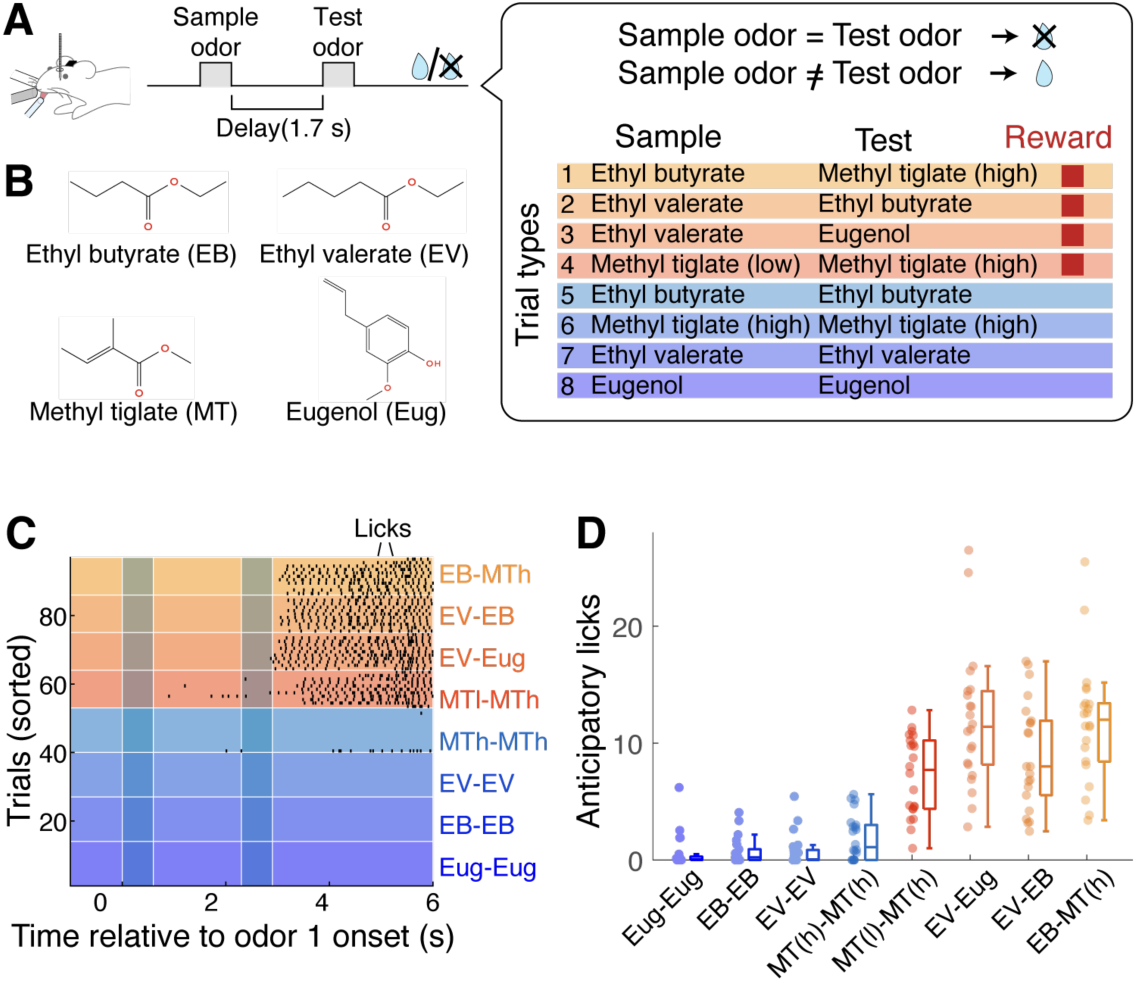
Head-fixed mice perform multiple-odor DNMS task proficiently. **A**, Experimental configuration for stimulus presentation and recording from head-fixed mice and trial structure. **B**, Molecular structures of odors used in the paradigm (left) and odor pairs for the 8 trial types (right panel). **C**, Lick raster from an example session from a trained mouse. Time zero corresponds to the onset of sample odor. Trials are sorted by trial types. **D**, Average number of licks during 3 s following the test odor presentation (“anticipatory licks”) for each trial type. Each data point corresponds to one mouse. N = 22 mice.

### PL and ORBl are most enriched in delay-period odor identity signals

To first distinguish how odor coding differs across frontal regions, we recorded single-unit activity acutely with Neuropixels 1.0 probes in head-fixed mice proficient in the multiple-odor DNMS task. We targeted both lateral and medial frontal regions, including the secondary motor cortex (MOs), frontal pole (FRP), agranular insular cortex (AId), lateral, ventrolateral, and medial orbitofrontal cortices (ORBl, ORBvl, ORBm), prelimbic cortex (PL) and the dorsal part of anterior cingulate cortex (ACAd). Across the 22 trained mice, 4742 units were classified initially by Kilosort 4.0 (Pachitariu *et al*, 2024) as good units. Of these, 2898 units were further manually sorted for waveform consistency, clean refractory period, and stability (Supplementary Fig. 2). Of these, 545 were registered to MOs, 158 to FRP, 204 to AId, 538 to ORBl, 501 to ORBvl, 470 to ORBm, 450 to PL, and 32 to ACAd (Fig. 2A).

Many units in multiple regions showed increased spiking activity during the delay period, consistent with the established literature that frontal cortical neurons can maintain sensory information (Fig. 2B). To quantify how much odor identity information each region carried during this period, we performed linear decoding analyses on population activity (Fig. 2C,D). For each region, ensembles of a given size were randomly selected (4–40 units; Fig. 2C), and used to train and decode the identity of the sample odor. Decoder accuracy increased with the ensemble size in all regions, indicating that delay-period activity contained non-redundant information about which sample odor had been presented.

**Figure 2:**
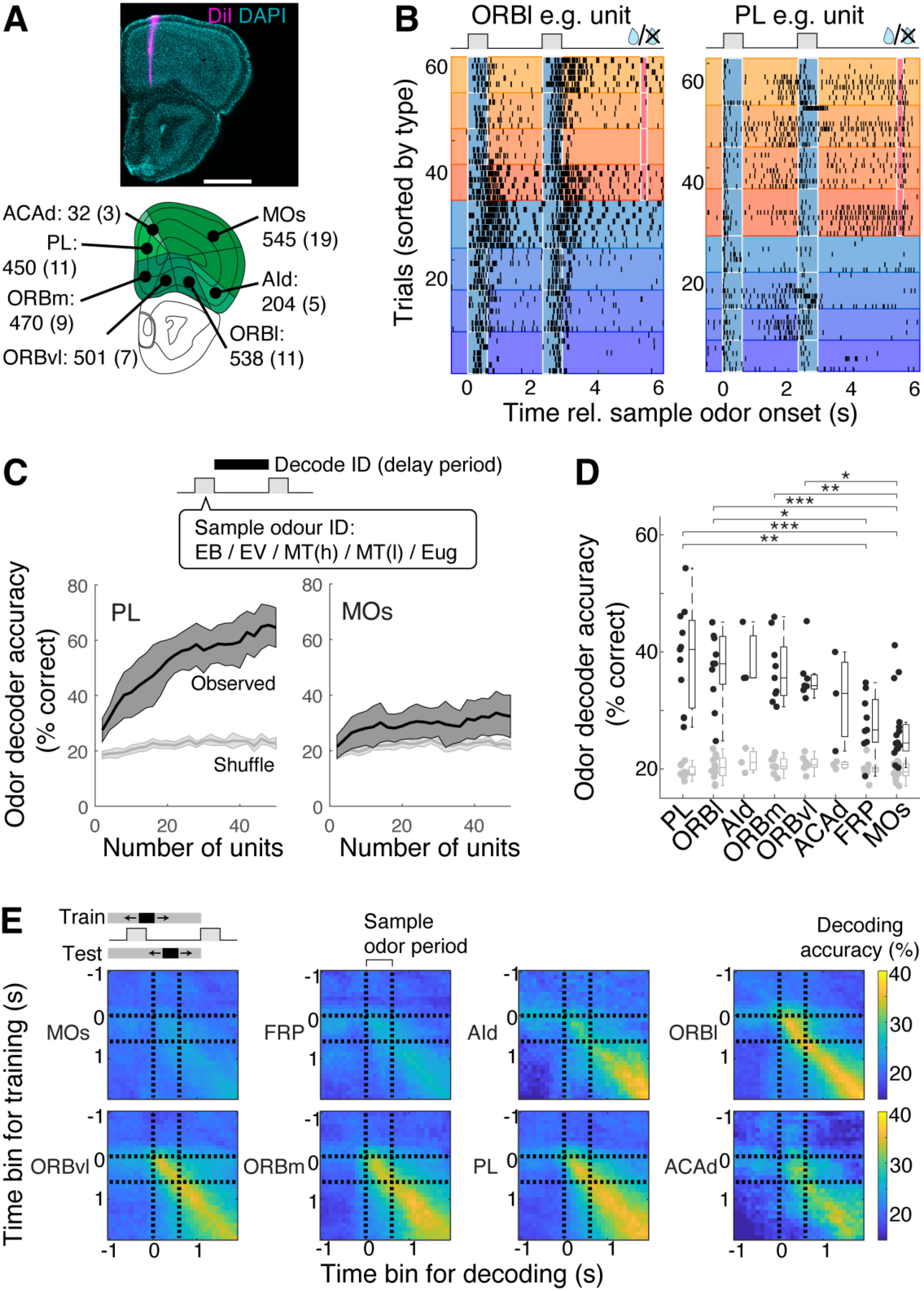
Regional variations in delay odor encoding. **A**, Top, Example confocal fluorescent image showing a coronal section stained with DAPI (cyan) at the level of frontal cortex and a track from a Neuropixel probe that had been coated with DiI (pink). Scale bar = 1 mm. Bottom, corresponding reference atlas (Allen) and the numbers registered units from all experiments. **B**, Example spike raster plots for an ORBl unit (left) and a PL unit (right). Trials are sorted in the same way as in Fig. 1C. Blue shaded period = odor presentation; red shaded period = water reward. **C**, SVM decoding accuracy to classify sample odor identities and its dependence on the number of units sampled from PL (left) and MOs (right). Delay activity was used for training and testing. Mean and S.D. shown. **D**, Accuracy of general odor identity decoding. Ensemble activity from 8 randomly sampled units used. Each data point corresponds to one mouse. Black and gray data correspond to observed and shuffled data, respectively. Dotted lines = onset & offset of sample odor. Regions were sorted by the median accuracy values. **E**, SVM decoding accuracy for the training time bins (rows) and testing time bins (columns) shown with the corresponding colormap.

Surprisingly, MOs carried the least sample odor information during the delay, even though the decoding performance was above chance (Fig. 2C,D; mean accuracy = 0.274 ± 0.061 and 0.284 ± 0.049 at 8 and 16 units; p = 0.002 and 0.0039 compared to shuffle control; Wilcoxon signed rank test; n = 14 and 10 mice, respectively). Decoder accuracy in MOs did not increase significantly with additional units (p = 0.7123, 1-way analysis of variance for the effect of sample size) and remained lower than other frontal areas for all ensemble sizes examined. In contrast, regions such as PL and ORBl exhibited the most accurate decoding of sample odor identity. For example, with the ensemble activity from 8 units, decoders trained on PL and ORBl data achieved the highest accuracies (mean accuracy = 0.402 ± 0.083 and 0.378 ± 0.064 for PL and ORBl, respectively; p = 0.002 and 0.001; Wilcoxon Signed rank test against shuffle control; n = 10 and 11 mice) and increased further with more units (p = 0.0002 and 0.0003, 1-way ANOVA for the effect of sample size). These results indicate that delay-period activity is markedly more informative about the sample odor identity in the medial and ventral lateral prefrontal regions such as PL and ORBl than in MOs.

We next asked whether delay-period representations are stable or evolve dynamically over time (Fig. 2E). To this end, we trained decoders on ensemble activity from a time bin that ranged from the sample odor onset until the end of the delay period, and tested the trained decoder on time bins that either matched or increasingly diverged from the training bin (Meyers *et al*, 2008). Decoder performance was highest when training and testing used the same time bin and decreased progressively as the temporal offset increased (Fig. 2E; Supplementary Fig. 3; Average time constant across brain areas = 4.455 ±1.768 s; n = 65 region-session pairs from 22 mice). This temporal specificity suggests that the delay-period population code is not static but drifts over time. However, even under this more stringent, time-resolved measure, PL maintained among the strongest odor identity signals across frontal regions (Peak accuracy estimate [α] for PL = 0.348 ± 0.070 at 8 units used; n = 10 mice). The results overall indicate that PL and ORBl reliably retain recent olfactory information, while MOs carries little sensory coding.

### Match vs.non-match choice signals are widespread but differ in stimulus invariance

Beyond maintaining sensory information during the delay period, match vs. non-match recognition and subsequent choice computation require that incoming test odor signals be gated, or modulated, by the sample odor. We therefore examined how activity evoked by the test odor was modulated by whether the sample and test odors matched. Many frontal neurons increased or decreased their firing selectively on match versus non-match trials, as illustrated by individual examples from MOs and PL (Fig. 3A,B).

**Figure 3:**
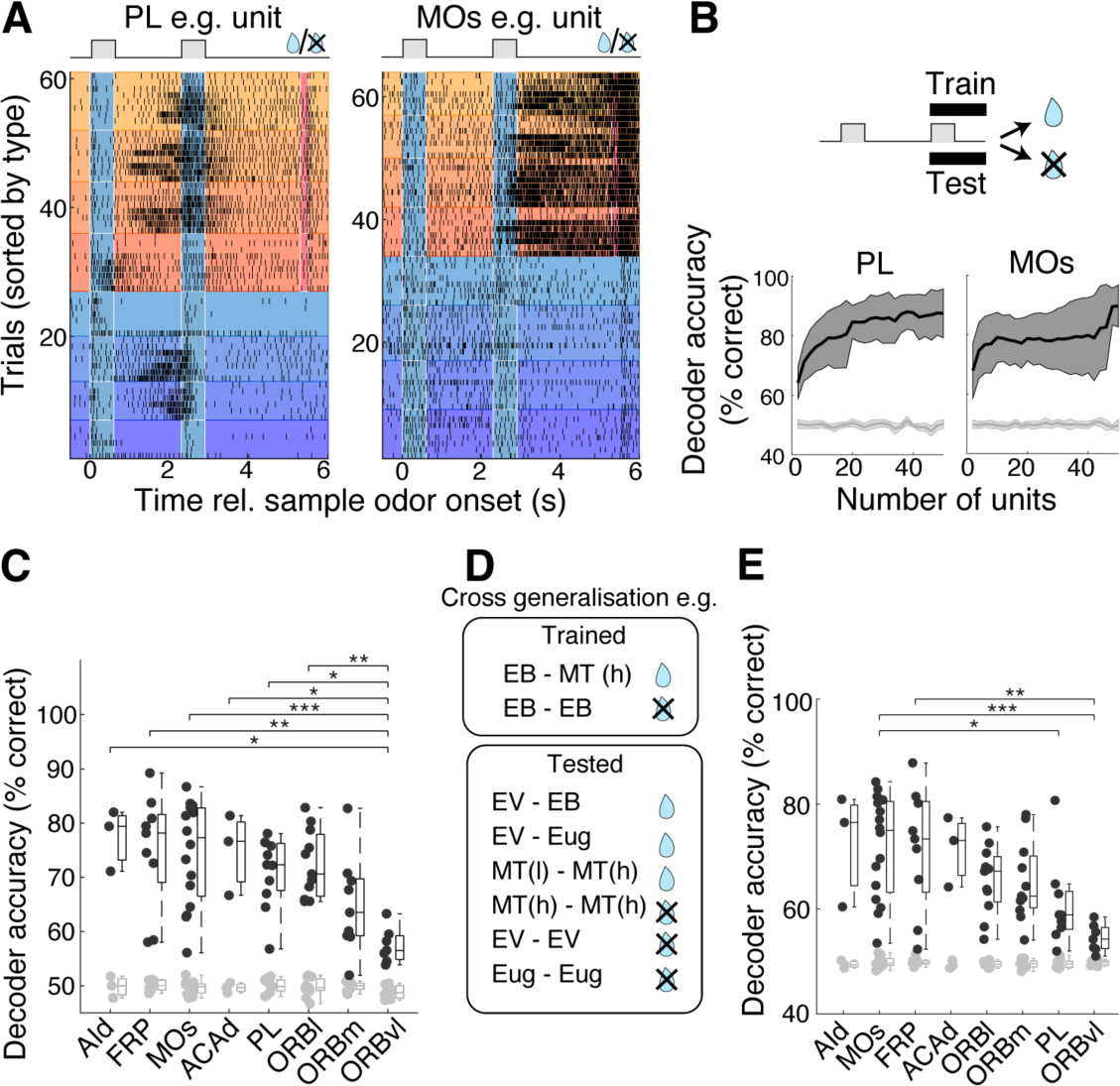
General vs. stimulus-invariant outcome encoding reveals regional differences. **A**, Example spike raster plots of a PL unit (left) and an MOs unit (right). **B**, Dependence of SVM decoding accuracy to classify the outcome and its dependence on the number of units sampled from PL (left) and MOs (right). Ensemble activity from 2 s following the test odor onset was used. Black and gray lines correspond to observed and shuffled data, respectively. Mean and S.D. shown. **C**, Accuracy of SVM outcome decoding using ensemble activity from 4 randomly sampled units. Black and gray points = observed and shuffled data. **D**, Cross-generalization of SVM decoding to reveal stimulus-invariant outcome encoding; trials with a specific sample odor were used for training (top panel), and the trained SVM was tested on other trial types (bottom panel) to decode the outcome. **E**, Accuracy of SVM cross-generalization. Ensemble activity from 4 sampled units used. Black and gray points = observed and shuffled data.

To quantify how robustly each region represented the match vs. non-match choice at the population level, we trained SVM decoders to classify the choice based on ensemble activity upon test odor presentation (Fig. 3B). For each region, given that the choice classification is binary, we randomly sampled a smaller group of neurons (4 units) and assessed how accurately the choice can be predicted from the test trial data. Choice decoding was high across most frontal regions, even with this low number of units (Fig. 3C; Average accuracy for all regions (auROC) = 0.710 ± 0.067; n = 65 region-session pairs from 22 mice). This suggests that reward-associated, choice-related information is widely distributed across the frontal cortex. One exception was ORBvl, which showed a comparatively poor performance (Average accuracy (auROC) = 0.574 ± 0.033, n = 7 mice; p = 0.0002, effect of brain regions in 1-way ANOVA; F = 4.86, degrees of freedom = 7)), indicating less enriched representation of choice.

We then asked whether these choice-related signals reflected a general match vs. non-match recognition, independent of specific odors or whether they were tied to a particular odor used. To dissociate these possibilities, we measured how the trained decoder cross-generalized across odor pairs (Fig. 3D,E). Specifically, we trained SVM to classify the choice using only trials with just one sample odor type and then tested its decoding accuracy on trials involving other odor pairs not used in the training set (Fig. 3D). If a region encodes a more abstract, stimulus-invariant match vs. non-match signal, a decoder trained on this restricted odor set should generalize well to other odors.

This analysis revealed a more varied decoding ability across the frontal subregions, where dorsolateral areas, including MOs, supported the strongest cross-generalization of match vs. non-match decoding across odor pairs (Fig. 3E; Average decoding accuracy for MOs (auROC) with 4 units = 0.733 ± 0.094, n = 16 mice). In contrast, as with the general outcome decoding, ventromedial regions such as ORBvl exhibited poor cross-generalization (Average decoding accuracy (auROC) with 8 units = 0.554 ± 0.025, n = 7 mice). Intermediate levels of cross-generalization were observed in anatomically intermediate regions such as PL and ORBl, which showed robust general choice decoding but only partial cross-generalization across odor pairs (Cross-generalising accuracy = 0.653 ± 0.060 and 0.643 ± 0.081 for PL and ORBl, respectively; n = 10 and 11 mice). Further, unlike MOs, the intermediate regions also exhibited above-chance levels of match vs. non-match decoding on error trials (Supplementary Fig. 4), rather than strictly predicting the behavioral outcome.

### Match-selectivity and early discrimination for outcome are enriched in the prelimbic area

Brain regions that participate in the intermediate stages of decision making may exhibit other activity patterns that reflect intermediary characteristics. For example, a model of match vs. non-match computation posits that a comparator circuit contains both match-selective and non-match-selective neurons (Engel & Wang, 2011). We therefore assessed which frontal cortical regions are enriched in neurons that show match selectivity, even though only non-match trials were rewarded. For each brain region, single unit firing rate following the test odor presentation was measured and the number of neurons that showed significant positive modulation for the unrewarded, match trials was counted. Only a small number of frontal regions contained match-selective neurons, and PL had the largest proportion (Fig. 4A,B). To analyse further how robustly the sample-test odor match is represented despite not being reinforced, a representational dissimilarity matrix was computed (Nili *et al*, 2014). This revealed that some regions, such as MOs and AId, exhibited consistent representation only within rewarded, non-match trials (Fig. 4C-F). On the other hand, intermediate region such as PL exhibited consistency in the ensemble activity for match trials in addition (Fig. 4C-E).

**Figure 4:**
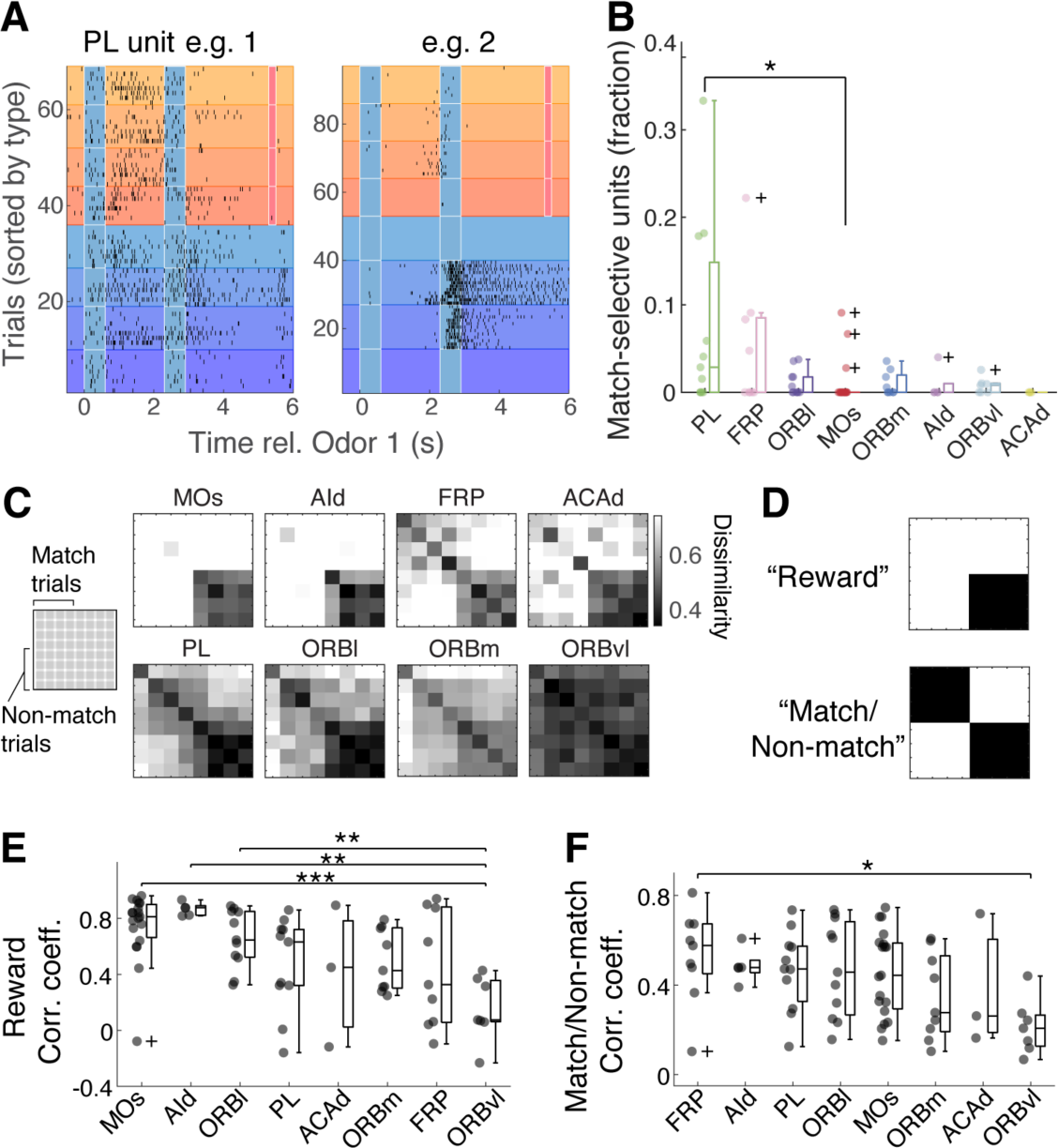
Intermediate stages of non-match computation. **A**, Example raster plots from two PL units that show match-selective firing rate increase. **B**, Numbers of units that significantly increase firing rate in response to test odor on match trials as proportions of total units registered to the region. **C**, Representational dissimilarity matrix (RDM) for 8 trial types, with gray level corresponding to dissimilarity (1 - correlation coefficient) averaged across mice. Match trials are sorted first, followed by non-match trials. **D**, Two models; top, activity patterns for rewarded trials are similar; bottom, activity patterns are similar within match, and non-match conditions. **E**, Correlation between region-specific RDM and the reward model shown in **D**. **F**, Correlation between region-specific RDM and the match/non-match model in D. ***, **, and * indicate significance at p<0.001, p<0.01, and p<0.05, respectively.

A further characteristic of an intermediary decision process may be temporal, with intermediate regions expected to show earlier choice modulation. To test this hypothesis, we quantified, for each neuron, the discrimination time between match and non-match trials. This is the time at which the firing rate of a given unit diverges significantly for non-match vs. match trials (Methods). Interestingly, the regions containing units with the earliest discrimination times were not the most generalized dorsolateral areas but instead overlapped with frontal regions that showed intermediate levels of cross-generalization, such as PL and ORBl (Supplementary Fig. 5).

To summarise and visualise the joint representation of sensory and choice signals, we computed UMAP embeddings of multiple decoding results for both variables (Fig. 5A-D). Sessions with high versus low behavioural accuracy occupied distinct regions of this embedding space (Fig. 5B), consistent with PFC population activity covarying with behavioural performance. Strikingly, under high-accuracy conditions, the low-dimensional embedding revealed a smooth medioventral-dorsolateral gradient (Fig. 5D), indicating that information about sensory inputs and choices is organised along this anatomical axis when the task is performed reliably. Taken together, these results support a spatial organisation in which the frontal cortex contains a distributed but structured arrangement for match vs. non-match computation. Ventromedial and orbitofrontal regions emphasize stimulus-specific information and encode outcome with limited generalization across odor pairs. Dorsolateral regions, including MOs, emphasize more abstract, stimulus-invariant match/non-match signals that generalize well across odors but carry relatively little sensory detail. Intermediate medial and lateral prefrontal areas such as PL and ORBl sit between these extremes, combining strong delay-period odor identity signals, robust outcome decoding, intermediate cross-generalization, and the earliest discrimination times. These characteristics may suggest their involvement in intermediary, comparator circuitry where choice-related modulation occurs in a stimulus-specific manner (Fig. 5E). In contrast, more abstract, stimulus-invariant decision representations may occur downstream through integration of these intermediary signals in a reward-dependent manner.

**Figure 5:**
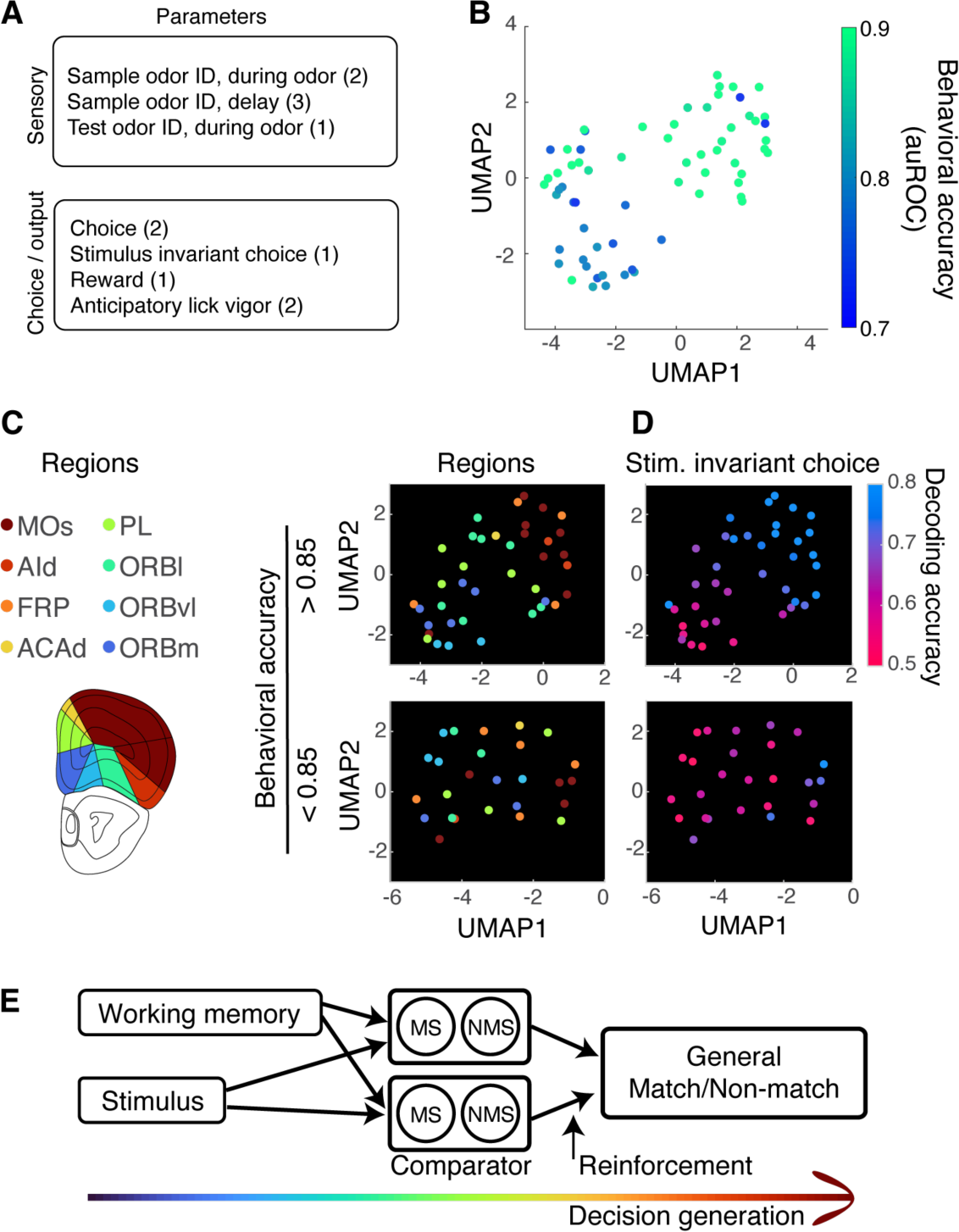
Gradual emergence of decision variables. **A,** Parameters extracted from decoding and regression analyses used for dimensionality reduction. Numbers in bracket correspond to the number of parameters in each category. **B**, UMAP embedding of the parameters listed in **A.** Individual points correspond to brain regions from individual behavioral sessions that contained enough numbers of units. The points are color-coded by mice’s accuracy in the multi-odor DNMS task. **C**, Left, Color codes corresponding to the brain regions. Right, UMAP embeddings of parameters in **A**, but with the data separated by the behavioral accuracy. Plots are color-coded by the brain regions. **D**, UMAP embeddings as in **C**, but color-coded by accuracy of decoding stimulus invariance choice (cross-generalization). **E**, Model of match vs. non-match computation based on Engel & Wang (2011), and possible hierarchy in the decision making corresponding to the gradual emergence of general outcome from sensory information. The intermediate circuitry compares the retained sensory information and incoming sensory information in a stimulus-specific manner. MS and NMS units correspond to match-selective and non-match-selective units. The detected match vs. non-match signals are integrated in a reward-dependent manner.

## Discussion

Our results reveal a functional gradient across the mouse frontal cortex during an olfactory working memory task. During the delay period, sensory information was sparse in dorsolateral regions such as MOs, whereas stimulus information was most enriched in ventromedial and intermediate areas. In contrast, choice-related activity was broadly distributed across PFC, except in ORBvl. However, the detailed nature of the choice signals differed across regions: Dorsolateral regions (MOs, AId) were dominated by stimulus-invariant, reward-associated choices, whereas intermediate areas (PL, ORBl) were enriched in stimulus-specific match/non-match activity with balanced selectivity for the two choices. This difference suggests regional differences in intermediate versus higher, more abstract stages of decision-making.

One key observation from our study is that MOs carried minimal information about sample odor identity when animals held this information in memory but a decision was yet to be made. This contrasts with robust sensory representations in MOs during a range of other acquired but more direct stimulus-response behaviors (Guo *et al*, 2014; International Brain *et al*, 2025; Le Merre *et al*, 2026; Steinmetz *et al*, 2019). Given that MOs receives input from orbital cortices (Harris *et al*., 2019; Heidbreder & Groenewegen, 2003; Hoover & Vertes, 2007), which are themselves connected with the olfactory areas and exhibit task-relevant, odor-driven responses (Critchley & Rolls, 1996; Ramus & Eichenbaum, 2000; Rolls *et al*, 1996; Schoenbaum & Eichenbaum, 1995a, b; Wu *et al*., 2020; Zhou *et al*, 2019), the absence of informative olfactory signals in MOs in our study is unlikely to be simply due to our choice of sensory modality. Instead, this pattern may reflect the difference in task structures: mnemonic processing may preferentially recruit mPFC regions such as PL, whereas MOs may be more engaged when sensory cues are coupled directly to action execution (Passingham, 2021).

The regional differences we observed align closely with known connectivity-based subdivisions of the frontal cortex, which comprise ventral, ventromedial, and dorsomedial divisions (Le Merre *et al*., 2021). Dorsomedial areas, such as MOs, are known for reciprocal connections with neocortical areas including the motor cortices (Harris *et al*., 2019; Le Merre *et al*., 2021; Li *et al*, 2020). This is consistent with the enrichment of general, choice-related activity in this region that we observed, as well as their involvement in a range of acquired, sensory-driven behaviors mentioned above. Ventral regions, on the other hand, receive strong inputs from sensory areas, including olfactory structures such as AON and piriform cortex. This is consistent with our observation that ORBvl and ORBm regions were informative about olfactory stimuli, but relatively less enriched in choice signals. Ventromedial areas, including PL, are characterized by dense local interconnectivity within PL, as well as afferents from ventral orbital areas and limbic areas (Harris *et al*., 2019; Hoover & Vertes, 2007; Le Merre *et al*., 2021). In turn, PL projects to dorsomedial regions such as ACAd and MOs (Heidbreder & Groenewegen, 2003; Le Merre *et al*., 2021; Sesack *et al*, 1989). This intermediate anatomical position is consistent with our functional characterisation, suggesting that PL occupies a transitional role in mnemonic decision-making.

It should be noted, however, that our descriptions are in terms of regional enrichments, rather than strict correspondence with anatomical boundaries. Functional organization does not map perfectly onto anatomical divisions, whether they are defined by cytoarchitecture or connectivity (Le Merre *et al*., 2026). Further, within each region, the functional signatures of neurons differ depending on the neurotransmitter and molecular profiles, as well as the neuron-specific connectivity patterns (Kamigaki & Dan, 2017; Kim *et al*, 2016; Ye *et al*, 2016). In this regard, the enrichment of match-selective neurons in PL may also reflect their involvement in inhibitory control of behavior (Courtin *et al*, 2014; Li *et al*., 2020), rather than serving as a reservoir of intermediate operations. Future work resolving cell-type-specific contributions will be critical for clarifying these mechanisms.

Often considered the apex of the cortical hierarchy, the prefrontal cortex is defined by dense reciprocal connectivity and widespread task- and choice-related activity. Through a rich yet tractable behavioral paradigm, our direct comparisons of subregions distinguished functional profiles aligned with different stages of decision-making. These findings provide a step toward understanding how mnemonic decisions are constructed and translated into behavior.

## Methods

### Animals

All protocols for animal experiments were approved by Animal Care and Use Committee of the Okinawa Institute of Science and Technology Graduate University [Protocol No. 2021-350]. Twenty-two adult male and female C57BL/6 mice (8 – 15 weeks old; Japan Clea) housed singly under a reversed 12-hour daylight cycle were used.

### Headplate Implantation

All recovery surgeries were performed under aseptic conditions. Animals were deeply anesthetized with isoflurane (3% induction, 1.5% maintenance; IsoFlo Zoetis, Tokyo, Japan) and placed on a stereotaxic frame (1900, Kopf Instruments, CA, USA). The animal’s temperature was kept at 36.5 °C using a heating blanket with a DC thermostat (40-90-8D, FHC, ME, USA). To attach a custom-designed aluminum headplate (1cm × 2.5cm × 0.2cm; ∼1g), the skull was exposed by gently removing the scalp and the underlying soft tissue. The bone was softly scarred using a dental drill (Success 40, Osada, Tokyo, Japan) and subsequently cleaned and dried. The stereotactic landmarks were marked for future electrode insertion (Lateral coordinate AP: +2.6, ML: +1.5; Medial coordinate: AP: +2.6, ML: +0.5) with a fine pen (Sakura Micron Pigma 005, Osaka, Japan). The skull was covered with cyanoacrylate (Histoacryl, B. Braun, Melsungen, Germany) before the headplate was fixed to the skull with gel super glue (Loctite, Henkel, Düsseldorf, Germany) and dental cement (Kulzer, Hanau, Germany). Mice were recovered for ∼2 hours in a warm chamber (37 °C) before being returned to their cages. Post-operative analgesic (Carprofen 5mg/kg, Rimadyl, Zoetis, Tokyo, Japan) was administered subcutaneously for 3 consecutive days.

### Olfactometry and Odors

For behavioral experiments, odors were delivered using a custom-made flow-dilution olfactometer described previously (Reuschenbach et al., 2023). Briefly, filtered and deodorized air (2 SLPM) was odorized by gating a separate stream of flow-controlled air through odor canisters to generate approximately 1% of saturated vapor. Sample and test odor were delivered through separate manifolds and Teflon-coated tubing to minimize cross-contamination, and the final timing of the odor presentation was controlled by actuating a 5-way solenoid valve. Inter-trial interval was approximately 40 seconds, during which the odor lines were purged with high-pressure air to minimize cross-contamination. Odors used were: ethyl butyrate (EB; #W242705; Sigma-Aldrich, MA, USA) and methyl tiglate (MT; #T0248; Tokyo Chemical Industry, Tokyo, Japan), ethyl valerate (EV; #290866; Sigma-Aldrich), methyl valerate (MV; #S0015; Tokyo Chemical Industry), methyl salicylate (MS; #V0005; Tokyo Chemical Industry), and eugenol (Eug; #A0232; Tokyo Chemical Industry).

### Behavioral training

#### Habituation

Habituation took place at least two weeks after the surgery. Water bottles were removed from the home cage in the morning of the first day of habituation. Over the next 3-5 days, the mice were head-fixed via the implanted head plate and placed on a custom-made running wheel. During this period, mice were trained to receive water by robustly licking from a water port in the behavioral setup, until they drank at least 1 ml of water. Animals’ licking pattern was recorded every time the infrared beam above the water port was broken and sensed (PM-F25 Panasonic, Tokyo, Japan). The respiration pattern was measured with a flow sensor (AWM3100V, Honeywell, NC, USA) placed in front of the animal’s right nostril.

#### Olfactory DNMS training

After habituation, mice were first trained on the two-odor version of the DNMS task with one odor pair (ethyl butyrate and methyl tiglate). Each trial start was indicated by a flashing blue LED signal (8 pulses at 8Hz) and was followed by a sample and test odor (0.6 s duration each), separated by a short delay period (1.7s). Non-matching trials were those where the sample and test odors were different. Mice were trained to associate a non-matching odor pair with a water reward (2 x 10 µl), delivered 2.5 s after the test odor offset. Trials were ordered in a semi-randomized order, such that no more than 3 consecutive trials were of the same rewarding condition. On average, 50% of the trials were non-match trials. The session ended when mice no longer generated licks for the water reward. Once mice reached 80% performance accuracy for both rewarded and unrewarded trials, they progressed to the next training stage.

#### Multiple-odor DNMS task

This DNMS task used five different olfactory stimuli (Ethyl butyrate, ethyl valerate, eugenol, and methyl tiglate at low and high concentrations (0.5% vs 1% of saturated vapor)). Odor pair combinations were selected to represent a range of perceptual similarities. Animals were exposed to non-match and match combinations in semi-randomized order, involving random permutation of the odor pairs to ensure that all combinations appeared, on average, in equal frequencies in each session. In subsequent sessions, the number of trial types was further reduced to four non-match and four match odor pair combinations. This reduction allowed a reasonable repetition of the respective trial types (i.e., odor pair combinations) in each session.

#### In vivo electrophysiology during Multiple Odor DNMS task performance

Neuropixels 1.0 probe ((Jun *et al*, 2017); IMEC, Leuven, Belgium) was connected to the acquisition system via an HS-1000 head stage. Signals were digitized using a PXIe-1000 acquisition card installed in a PXIe-1082DC chassis (National Instruments, Emerson, MO USA). Channels were synchronized via externally generated TTL signals with random temporal jitters with a mean inter-pulse interval of 0.84 ± 0.007 s, delivered from a multi-function data acquisition device (USB-6212, National Instruments, TX, USA). These TTL pulses were also used for aligning the electrophysiology data with the behavioral data. Neural signals were sampled from all 384 active recording sites at 30 kHz using the Open Ephys GUI (Siegle *et al*, 2017). The internal probe tip served as a reference. When required, probes were sharpened prior to experiments using a micropipette grinder (EG-45 Narishige, Tokyo, Japan) according to the procedure described previously (Steinmetz *et al*, 2021).

On the day of the recording, mice were anesthetized with isoflurane (2% induction, 1% maintenance; IsoFlo Zoetis), and a craniotomy was performed over the marked target site. Isoflurane was removed, and the mice were head-fixed in the recording setup and allowed to recover on a heating pad. The Neuropixel 1.0 probe, pre-coated with DiI (Thermo Fisher Scientific, MA, USA), was inserted using a micromanipulator (SM-8, Luigs & Neumann, Ratingen, Germany). The dura was pierced with a fine needle to facilitate the probe insertion. Neural spike activity was monitored during probe advancement, and the final depth was adjusted as needed (DV ∼2.9mm relative to the brain surface). The craniotomy was covered with 4% agarose (A9539, Sigma-Aldrich) in Ringer’s solution (NaCl (135 mM), KCl (5.4 mM), HEPES (5 mM), MgCl_2_ (1 mM), CaCl_2_ (1.8 mM) with pH adjusted to 7.3) for stability. Recordings began after at least 15 minutes of recovery from isoflurane anesthesia.

#### Post-hoc histology

After electrophysiological recordings, mice were deeply anesthetized with isoflurane and transcardially perfused with phosphate buffer (NaH_2_PO_4_ (225.7 mM), Na_2_HPO_4_ (774 mM), with pH 7.4), followed by 4% formaldehyde dissolved in the phosphate buffer. Brain tissues were dissected and post-fixed for ∼24hrs at 4 °C. The tissues were then sectioned at 100µm using a vibratome (Campden 5100mz-Plus, Loughborough, UK). Sections were stained with DAPI, and probe tracks were verified by imaging DiI fluorescence using a Zeiss LSM 900 confocal microscope with a 10x dry objective (10x/0.3 Plan-Apochomat WD2.0; Oberkochen, Germany).

### Data Analysis

#### Behavioral data analysis

Behavioral data were acquired at 1 kHz with a data acquisition device (USB-6343, National Instruments) and imported into Spike2 to detect event times for trial-start LED presentations, odor valve opening, water valve opening, and lick onsets and to extract the record of trial types. The event times were further analysed in MATLAB (2024a, Mathworks) using custom scripts.

Accuracy of DNMS task performance was quantified by comparing the number of anticipatory licks generated for rewarded vs. unrewarded trials and quantifying the discriminability using the area under the receiver operating characteristic curve (auROC). AuROC values were calculated in blocks of 50 trials using the MATLAB function *perfcurve*. Values of 0.5 indicate chance performance, whereas values approaching 1 indicate high accuracy. The progression of performance over trials was visualized using logistic regression implemented with the MATLAB function *glmfit*.

#### Spike Sorting

High-pass filtered analog data and spike-sorted using Kilosort 4.0 (Pachitariu *et al*., 2024) with modified default parameters and the appropriate channel map for the Neuropixel 1.0 probe. Sorting output was manually reviewed in Phy (Rossant *et al*, 2016). Good units were those clusters with a clean refractory period, well-defined waveforms distinct from noise, and stable amplitudes throughout the recording.

#### Anatomical alignment of recording sites

The Neuropixel probe tracks were reconstructed from confocal fluorescent images of coronal slices using the Neuropixel trajectory explorer (https://github.com/petersaj/neuropixels_trajectory_explorer) allows for manual adjustment of the probe position within the Allen CCFv3 (Wang *et al*, 2020) to assign brain regions along the depths of the probe. This alignment was cross-validated using HERBS toolbox (Fuglstad *et al*, 2023). The final alignment was based on the Neuropixel trajectory explorer output since the two methods gave comparable results with minor differences. Each unit was assigned to a registered brain region according to its depth and channel location along the probe shank.

#### Linear decoder analysis for odor identity

Support vector machine (SVM) was trained with average firing rates during the late phase of sample-test odor delay period (1 second window before the test odor onset) of 4-40 randomly selected units from each brain region (“ensemble activity) to decode which of the 5 olfactory stimuli were presented during the sample odor. 80% of randomly selected trials from the behavioral session were used for training and the remaining 20% were used for testing the decoding accuracy. Accuracy is the % of the trials where the decoded odor was the same as the presented odor. This procedure was repeated 100 times for each ensemble size, and the decoding performance was averaged. Shuffle control was obtained by randomly permutating the trial types using the Matlab function *randperm*. Sessions were excluded from specific regional analyses if insufficient units were registered to the brain region concerned.

#### SVM for choice decoding

For each run, a subset of units was selected for the decoding analysis randomly, without replacement. The ensemble activity from 80% of randomly selected trials were used to train SVM to decode whether the trials were rewarded (non-match) or unrewarded (match). Random sampling was implemented using the MATLAB function *randsample*. The remaining 20% of the trials were used to test the decoding performance. MATLAB function *fitcecoc* was used to train the decoder with the Prior parameter set to uniform, and the decoding performance was tested using the MATLAB function *predict*. Ensemble sizes were varied from 4 to 40 and repeated 100 times each.

#### Cross-generalization SVM

This procedure involved training SVM on a specific sample odor (“A”) and testing the performance on the rest of the sample odors (“B – E”). With our 8 trial types, this was conducted in two implementations. The first training set was where the sample odor was ethyl butyrate (ethyl butyrate – methyl tiglate (high) and ethyl butyrate – ethyl butyrate pairs), and the testing sets comprised all other trials. The procedure was repeated for the training set where ethyl valerate was the sample odor. The overall decoding performance was the average from these two conditions.

#### Dynamic coding

SVM was trained with ensemble activity from 8 randomly selected units to decode the identity of the sample odor, where 80% of the trials were used for training and 20% for testing. The 0.5 s time bins for training and testing were systematically shifted covering the period from 1 second preceding the sample odor onset until the end of the inter-odor delay period. The decoding performance for each pair of training and test time bins were collected into a 2-dimensional cross-temporal decoding matrix.

#### Representational dissimilarity

To measure the dissimilarity of ensemble activity across trials in region, Pearson’s correlation coefficient was measured and subtracted from 1. To visualize the result, the pairwise distances were organized in the Representational Dissimilarity Matrix (RDM), which was a square matrix containing the above pair-wise dissimilarity with the trials sorted by trial type (same order as presented in Figure 1). To compare how much the trial-by-trial dissimilarity is explained by reward association, the pairwise ensemble activity dissimilarity was compared against the reinforcement model, where correlation value was 1 within rewarded trials but 0 across trial types, as well as within unrewarded trials. For the match/non-match selective model, the pairwise correlation was 1 within the same reward class for both match and non-match trials, but 0 across classes.

#### UMAP embedding

To represent the regional activity patterns in low dimensions using UMAP embeddings, the following parameters were extracted from Neuropixel recordings: accuracy of decoding the identity of sample odor during the odor presentation (1 s window from the onset) as well as the delay period (1 s window immediately following the sample odor offset; 1 s window preceding the onset of test odor, and the entire delay duration); accuracy of decoding the identity of test odor (1 s window from the onset); accuracy of decoding the choice following the test odor presentation (1 s window from the onset; 1 s window preceding the onset of water reward time; and 3 s window from the onset of test odor); decoding outcome (1 s window from the onset of water delivery time); cross-generalization decoding accuracy of choice where the training was limited to specific sample odor (using 1 s and 3 s windows from the test odor onset); correlation between representational dissimilarity matrix following test odor presentation and the reward outcome; correlation between predicted number of anticipatory licks and observed licks, based on ensemble activity following the test odor presentation (1 s and 3 s windows). Ensembles of 4 units were randomly sampled for each region for each instance of decoding and regression analysis, which was repeated 100 times for each condition. The parameters from each region and session were embedded in Uniform Manifold Approximation and Projection (UMAP) using the MATLAB (2026a) algorithm *umap* with the number of dimensions set to 2, and the number of nearest neighbors set to 10.

## Acknowledgement

We thank Yu-Pei Huang and Sayori Gordon for technical and administrative assistance, Josh Siegel for the support in setting up the Neuropixel data acquisition, and OIST’s core facilities (Animal Resources, Scientific Imaging, and Engineering Sections). We thank Japan Society for the Promotion of Science (JR, 202410751) and OIST Graduate University for the funding.

## Supplementary Figures

**Supplementary Figure 1:**
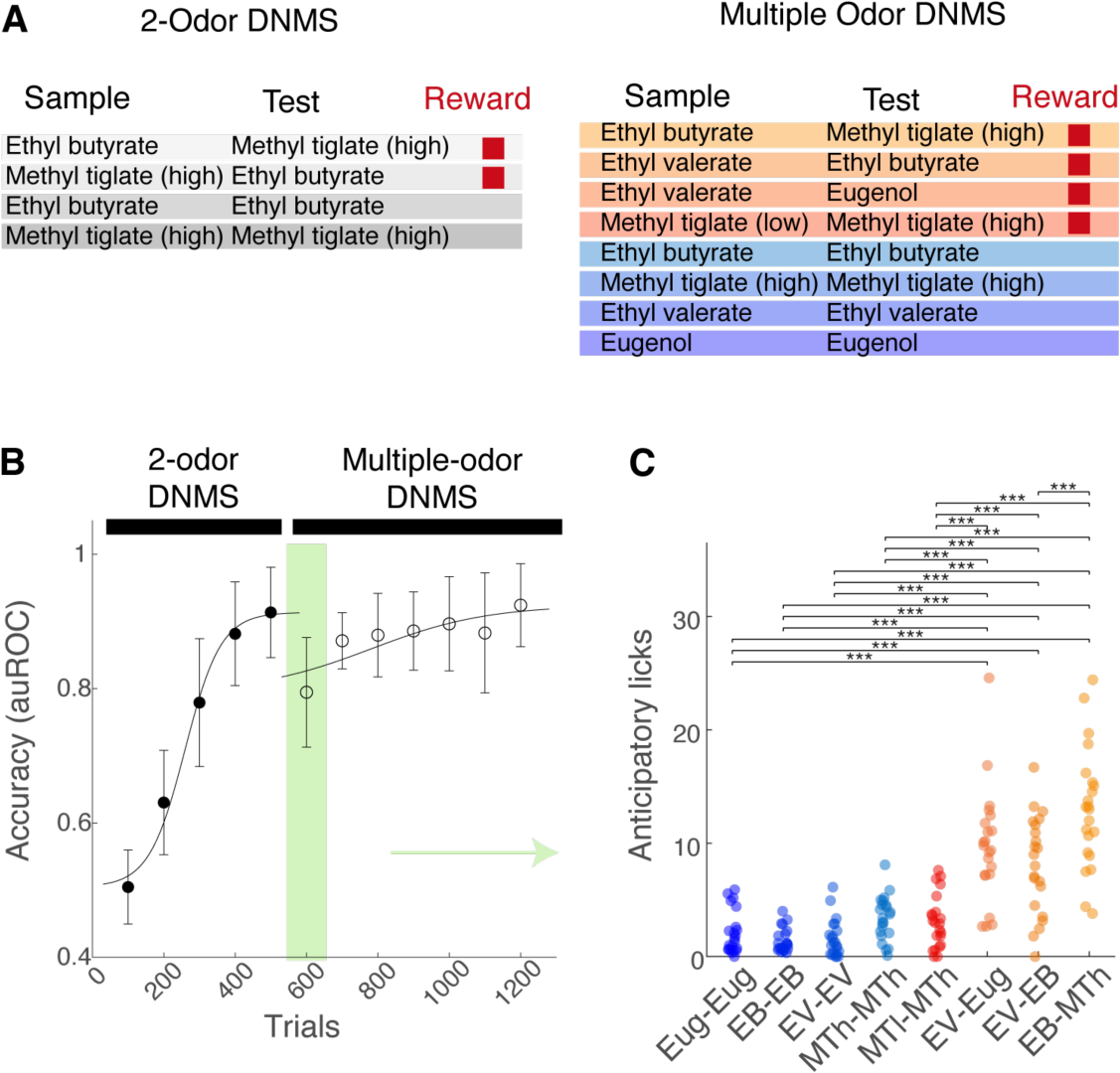
Behavioral readout of graded perceptual similarity **A**, Task rules for the 2-odor DNMS paradigm (left) and multiple-odor DNMS paradigm (right). **B**, After becoming proficient at the 2-odor DNMS paradigm, the DNMS task rule was generalized to multiple odors, ranging in the perceptual similarity. **C**, Behavioral accuracy on the first session after switching to the multiple-odor DNMS training. The mice showed above-chance overall performance here, indicating a robust transfer of the task rule. However, the performance depended on the odor pairs used, indicating a range of perceptual similarity between odor pairs.

**Supplementary Figure 2:**
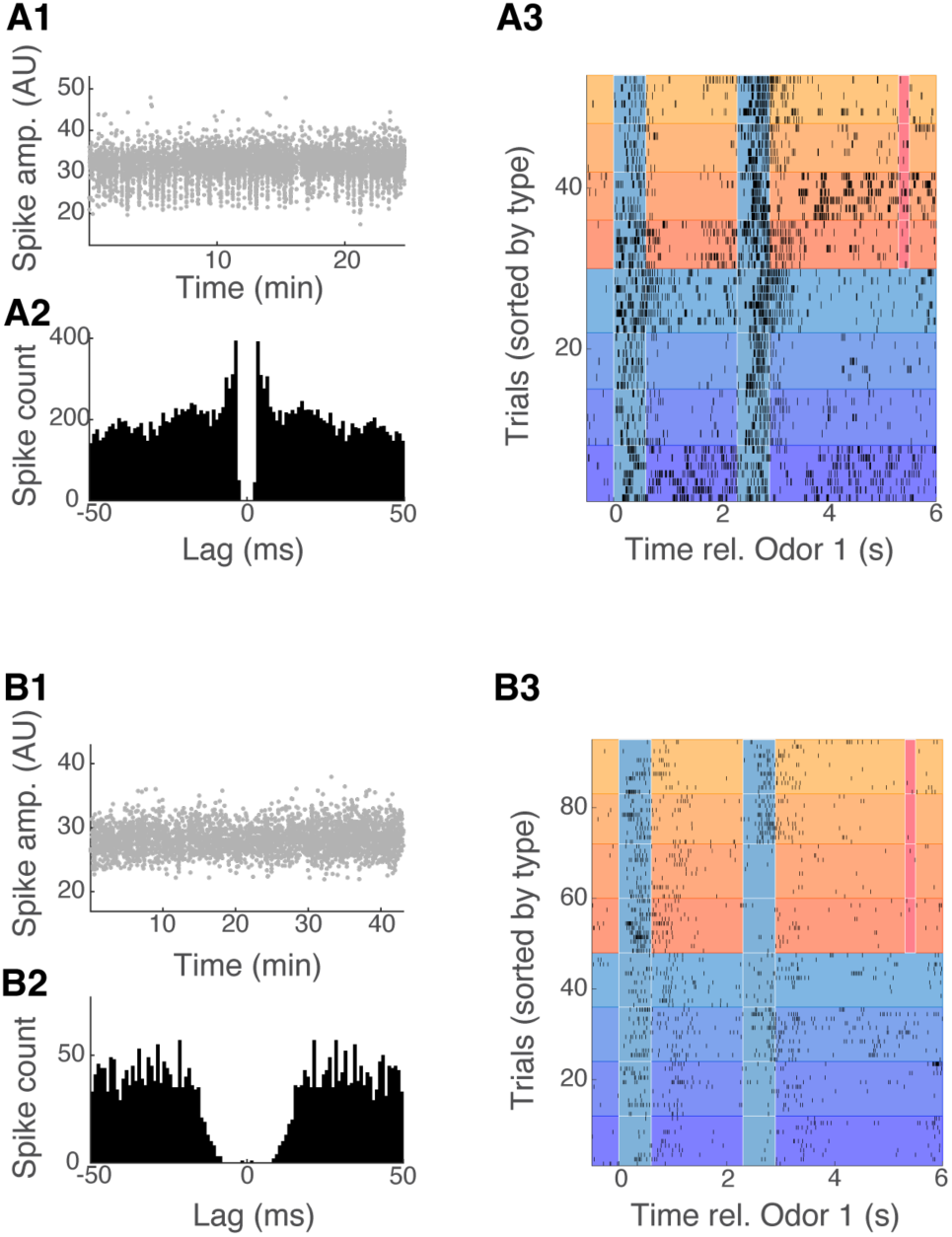
Examples of well isolated, stable units **A1-3**, Example unit 1. Units were first sorted using Kilosort 4.0. Of the units labelled “good” by the Kilosort algorithm, only those with stable spike amplitude (A1) and a clean refractory period based on auto correlogram (A2) were admitted for further analyses. Spike raster plot for this example unit during multiple-odor DNMS. **B1-3**, same as A1-3, but for another example unit.

**Supplementary Figure 3:**
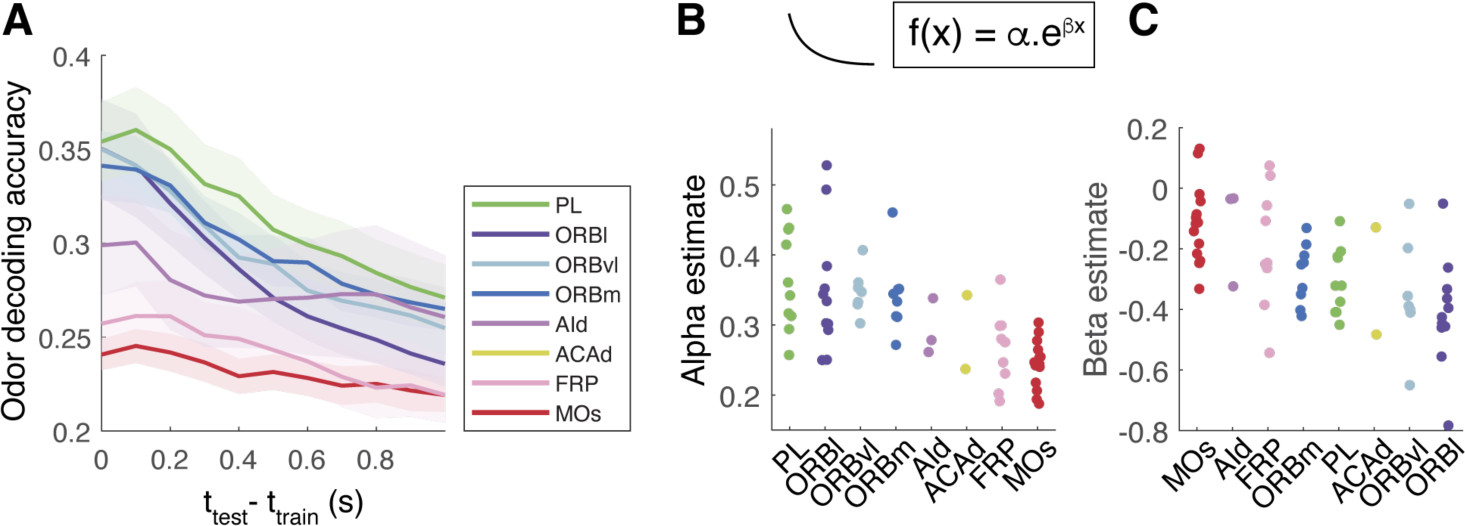
Time course of dynamic olfactory encoding during delay **A**, Odor decoding accuracy is highest when the time bins for training and testing are close together. **B**, **C**, The time course of dynamic coding was quantified by fitting the decoding accuracy curve over the temporal deviation (|t_test_ – t_train_|) by a single exponential function, and fitted coefficients α and β are plotted in panels B and C.

**Supplementary Figure 4:**
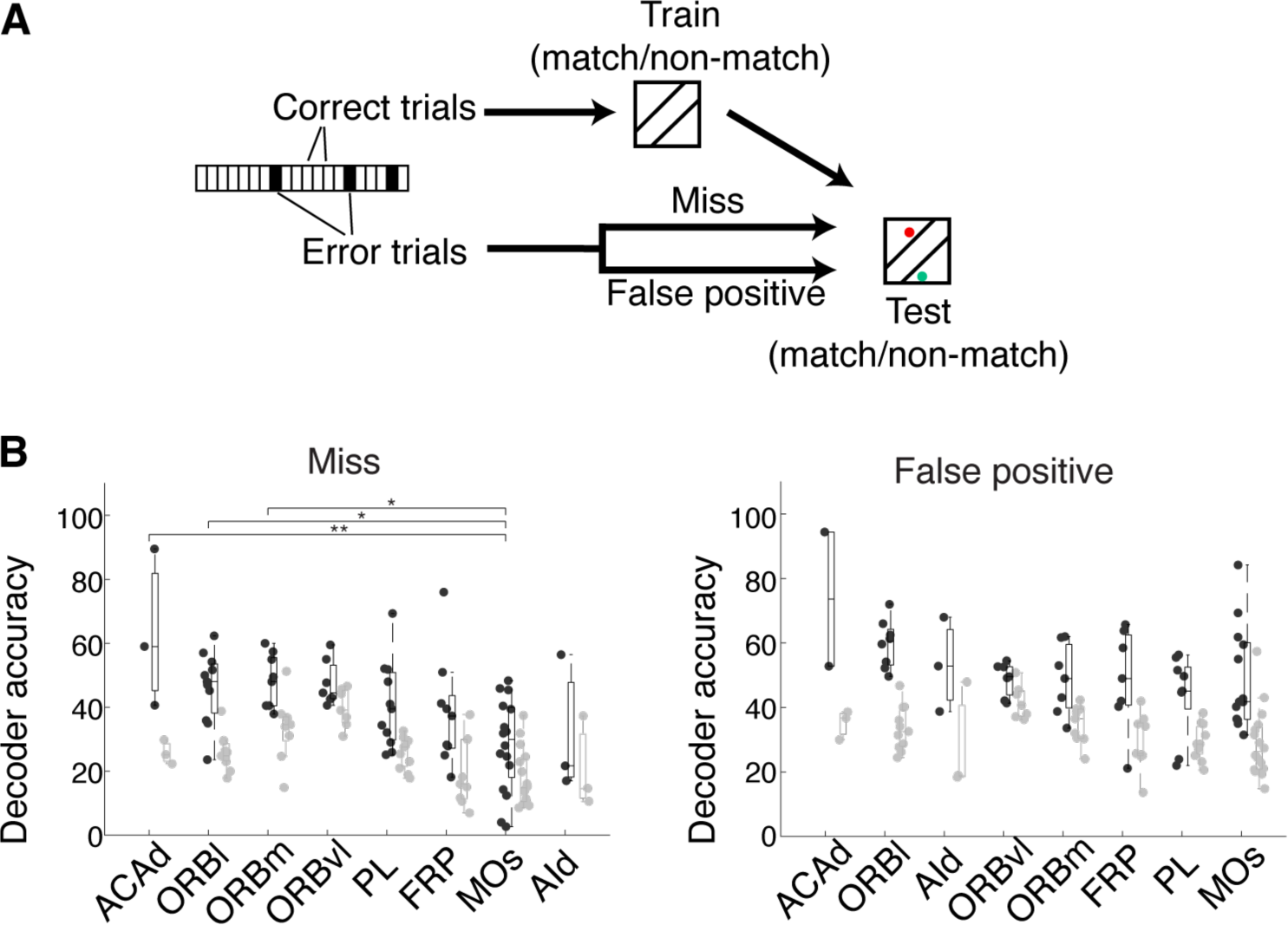
Error analysis suggests MOs encodes reinforced behavioral output **A**, SVM was trained to classify whether the choice period activity reflected match (unrewarded) vs. non-match (rewarded). Ensemble activity from 4 randomly selected units following the test odor presentation on correct trials was used. The trained SVM was tested on activity from error trials. “Miss” trials, where the mice failed to generate anticipatory licks on rewarded (non-match) trials were analyzed separately from “False positive” trials, where anticipatory licks were generated on unrewarded (match) trials. **B**, Decoder performance for the “miss” trials (left) vs. “false positive” trials (right). **, and * indicate significance at p<0.01 and p<0.05, respectively, with multiple test following a 1-way ANOVA to assess the dependence on brain regions.

**Supplementary Figure 5:**
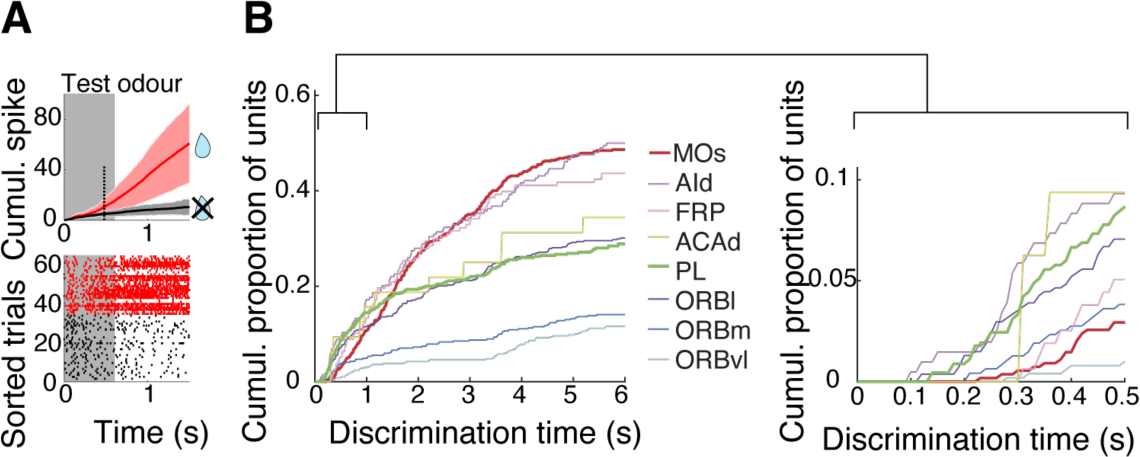
Intermediate regions show early divergence for the outcomes **A**, Top, for each unit, the discrimination time was defined as the earliest time at which the cumulative spike counts between rewarded vs. unrewarded trials significantly diverge after the onset of test odor according to Student’s t-test (p = 0.05). Mean and standard deviation for the two outcome types shown for this example unit. Bottom, Spike raster plots for the same unit, where spike times for the rewarded trials (red) and unrewarded trials (black) are shown. **B**, Cumulative proportion of units with the indicated discrimination times for the frontal regions as color coded. Inset (right) shows the first 500 ms expanded.

